# A Novel *Sarocladium spinificis* Strain Suppresses *Coccidioides posadasii* Growth: Morphological and Genetic Perspectives

**DOI:** 10.1101/2025.03.31.646452

**Authors:** Marcus de M. Teixeira, Sarah A. Ahmed, Heather Mead, Sybren de Hoog, Nathan Wiederhold, Jason E. Stajich, Daniel R. Matute, George Thompson, Bridget M. Barker

**Affiliations:** Núcleo de Medicina Tropical, Faculdade de Medicina, Universidade de Brasília, Brasília, Brazil; The Pathogen and Microbiome Institute, Northern Arizona University, Flagstaff, Arizona, USA; Radboudumc-CWZ Centre of Expertise for Mycology, Nijmegen, The Netherlands; Foundation Atlas of Clinical Fungi, Hilversum, The Netherlands; Translational Genomics Research Institute, Flagstaff, Arizona, USA; University of Texas Health Science Center at San Antonio, San Antonio-TX, USA; Department of Microbiology & Plant Pathology and Institute for Integrative Genome Biology, University of California, Riverside, Riverside, CA USA; Department of Biology, University of North Carolina at Chapel Hill, USA; Department of Medicine/Division of Infectious Diseases and Medical Microbiology and Immunology, University of California, Davis, CA, USA

**Keywords:** *Sarocladium spinificis*, fungal pathogens, comparative genomics, opportunistic infections, *Coccidioides posadasii*, antifungal resistance, microbial interactions

## Abstract

*Sarocladium* is a diverse fungal genus with implications in both plant and human diseases, exhibiting roles in pathogenicity, endophytism, and biocontrol. This study characterizes two *Sarocladium spinificis* strains, CA16 (CBS 144516) and CA18 (CBS 144517), isolated from patients initially suspected of having coccidioidomycosis. Morphological, phylogenetic, and genomic analyses confirmed their classification within the *Sarocladium* genus, closely related to *S. spinificis*. Both strains exhibited growth at 24°C and 37°C, with distinct morphological features. Comparative genomic analysis identified unique orthologous clusters and functional gene families associated with adaptability and potential pathogenicity. Notably, these strains demonstrated enhanced production of secreted proteins, CAZymes, and peptidases, highlighting metabolic versatility. Co-cultivation experiments revealed that *S. spinificis* strains CA16 and CA18 inhibited the growth of *Coccidioides posadasii*, suggesting competitive interactions and potential antifungal properties. These findings underscore the ecological and clinical significance of *S. spinificis* and its impact on microbial communities, advancing our understanding of its diversity, pathogenic potential, and antifungal capabilities.

**Importance:** Accurate identification and characterization of emerging fungal pathogens are essential for improving diagnosis and treatment. *Sarocladium spinificis*, though not widely recognized as a human pathogen, exhibits traits suggesting a potential role in opportunistic infections. This study provides the first detailed morphological, phylogenetic, and genomic characterization of clinical *S. spinificis* isolates, highlighting its ability to survive at body temperature and its antifungal resistance. Additionally, we demonstrate that these strains inhibit *Coccidioides posadasii*, the agent of Valley fever, suggesting ecological competition or antifungal properties. These findings contribute to understanding fungal interactions in clinical and environmental settings, with implications for fungal pathogenesis and antifungal strategies. By uncovering new aspects of *S. spinificis* biology, this work expands knowledge of *Sarocladium* species in human health and lays the foundation for future research on its ecological role, pathogenic potential, and therapeutic applications.

## Background

The genus *Sarocladium* was formally recognized in 2015, initially comprising 10 species, and has since gained attention for its diverse ecological roles and emerging clinical relevance. (1). The taxonomic recognition of *Sarocladium* as a new fungal genus was initiated by Gams and Hawksworth in 1975, acknowledging its implication in the manifestation of sheath-rot in rice. (2). This genus was initially characterized as *Acremonium*, but after a careful morphological examination, *Sarocladium* and *Acremonium* can be distinguished based on distinctive microscopic characteristics and multi locus sequencing type methods (3, 4). *Sarocladium* exhibits elongated phialides emerging individually on vegetative hyphae or on conidiophores with sparse or repeated branching, abundant adelophialides, and elongated conidia. In contrast, *Acremonium* typically features mainly unbranched or poorly basitonously branched conidiophores, variable-shaped conidia (subglobose, obovate, ellipsoidal), and the usual absence of adelophialides (1). With the advances of phylogenetic species concepts in fungi, multi-locus sequencing typing revealed that *Sarocladium* spp., *Parasarocladium* spp., *Chlamydocillium* spp. and *Polyphialocladium* sp. form a unique family within Hypocreales, named Sarocladiaceae (4, 5). *S. oryzae* and *S. attenuatum* were designated as type species of *Sarocladium* sharing ability to causing sheath rot in rice plants (1). The genus *Sarocladium* encompasses a diverse array of ecological functions, with (6) recognizing at least 37 species within its classification. These species play diverse roles, from saprophytes decomposing organic matter to pathogens infecting plants and humans. They also form endophytic associations, engage in mycoparasitism, and influence soil ecosystem dynamics (3, 7, 8). *Sarocladium* has gained recognition as a biocontrol agent against various plant diseases due to its ability to produce secondary metabolites that suppress fungal growth (9–11).

Contrastingly, *Sarocladium* spp. are abundant commensal fungi in the human mycobiome. Patients diagnosed with ulcerative colitis had lower levels of *Saccharomyces* and *Sarocladium* taxa and increased numbers of *Candida* spp. compared to healthy controls (12). Notably, *Sarocladium* infections in human are on the rise. At least 309 cases of human infections caused by *Acremonium* and *Sarocladium* have been described so far (8). *Sarocladium* species, including *Sarocladium kiliense*, *S. hominis*, *S. bactrocephalum*, *S. bifurcatum*, *S. pseudostrictum*, *S. subulatum*, and *S. terricola*, have been identified as potential causative agents of opportunistic infections in humans (1, 8). Mostly those isolates are able to grow at 37 °C, a characteristic shared by many of the human fungal pathogens (1, 4). Meningitis associated with *Sarocladium* sp. have been recurrently described in patients with immunological disorders (13–16). An outbreak of bloodstream infections attributed to *S. kiliense* emerged following the administration of contaminated antinausea medication in Colombia and Chile between 2013 and 2014. Through comprehensive whole-genome analysis, investigators identified a shared source of infection, confirming the presence of the fungus in contaminated vials of the medication (17). Lastly, recent reports have identified *Sarocladium* infections in individuals diagnosed with COVID-19 (18, 19). Understanding the implications of opportunist and concurrent infections, such as those involving *Sarocladium*, is crucial for healthcare professionals to ensure comprehensive diagnosis and treatment strategies.

To uncover the fungal etiology behind suspected coccidioidomycosis cases, our study yielded a surprising revelation: the isolation of two *Sarocladium spinifis* strains. Delving deeper, we undertook comprehensive phylogenetic and genomic analysis as well as co-cultivation experiments with *Coccidioides*. Comparative genomic analysis reveals unique genomic traits in *Sarocladium* species causing human infections. Remarkably, our findings demonstrated the suppressive effect of *Sarocladium spinifis* on *Coccidioides*, shedding new light on potential avenues for antifungal treatment.

## Materials and Methods

### 1 - Fungal isolation and phenotypical characterization

*Sarocladium* sp. isolates CA16 and CA18 were recovered at the Fungus Testing Laboratory at the University of Texas Health San Antonio, TX. The strain CA16 was obtained from a skin (lip) lesion while CA18 was isolated from a left leg wound, both in Madera, California, USA. These isolates were originally derived from ACCUPROBE-confirmed cases of coccidioidomycosis and were subjected to phylogenomic analysis of *Coccidioides*. Monosporic isolation procedures were conducted within a Biosafety Level 3 (BSL-3) facility at Northern Arizona University due to the presumed risk posed by these fungi classified as risk group 3. Mycelia were cultivated on 2X GYE (1% glucose, 2% yeast extract) agar at 25°C for a duration of 10 days. Macroscopic morphology assessment involved inoculating the fungus on various media including 2% Malt Extract Agar (MEA, manufacturer), Potato Dextrose Agar (PDA), Oatmeal Agar (OA), and 2% Glucose-1% Yeast Extract (2X GYE) plates, which were then incubated for seven days at 25°C. Additionally, a 2X GYE plates were incubated at 37°C for seven days, and morphology was evaluated for all media types. For microscopic characterization, the slide culture method was employed, wherein mycelia from macrocolonies were inoculated into blocks of 2X GYE, MEA, PDA, and OA agar. These blocks were covered with glass coverslips and incubated for one week. After growth, the coverslips were mounted in lactic acid and visualized with Zeiss EVOS® FL Auto Imaging System (Life Technologies) and micrographs were taken.

The competition assay between *C. posadasii* Δcts2/Δard1/Δcts3 (NR-166), which can be handled in a biosafety level 2 (BSL-2) and *S. spinificis* CA16 and CA18 strains, were performed on 2X-GYE a petri dish at 25°C since both species grow well on those conditions. We harvested conidia cells of both *C. posadasii* and *S. spinificis* using protocols described in (20). Next, we incoulated 10^7^ *C. posadasii* NR-166 conidia cells and uniformly spread onto 2X-GYE plates. We let the plates dry and 10^6^ conidia cells of *S. spinificis* CA16 strain or CA18 were plated in the center of the petri dish. We incubated the plates in the dark at 25°C for 10 days and captured images using an Olympus SZ-11 camera.

To characterize the susceptibility of the two *S. spinificis* isolates to antifungal drugs was conducted upon receipt of the samples using the broth macrodilution method, following the CLSI M38-A2 reference guidelines (21, 22). Minimum inhibitory concentrations (MICs) were determined as the lowest antifungal concentration that achieved an 80% reduction in fungal growth compared to drug-free controls.

### DNA extraction and molecular characterization

We extracted high molecular weight DNA from approximately 500 mg of freshly harvested mycelial tissue of the two *S. spinificis* strains, obtained seven days post-inoculation on 2X GYE plates. For DNA isolation, we employed the UltraClean® Microbial DNA Isolation Kit (MoBio, Qiagen). Fungal cell lysis was achieved through vigorous agitation in MicroBead tubes from the kit, using a vortex adapter tube holder (MoBio, Qiagen) for 15 minutes. To assess DNA quality and quantity, we performed electrophoresis on a 0.8% agarose gel stained with SYBR Safe (Thermo Fisher Scientific) and measured absorbance using a NanoDrop® ND-1000 system (Thermo Fisher Scientific). Given the initial diagnosis of the isolates as *Coccidioides* spp., we used a real-time-PCR based test targeting a specific repetitive region unique to *Coccidioides* genome, which yielded negative results (23). We used universal rDNA barcodes to differentiate between *Coccidioides* and *Sarocladium* (24). Approximately 20 ng of DNA served as input for the PCR amplifications of the ITS region as follows: 25µL of MyFi™ Mix (Bioline), 0.2 µM of each primer, DNA template and ultrapure water to a final volume of 50µL. The thermocycling steps were: initial denaturation for 2 min at 95°C, followed by 35 cycles of 15 s at 95°C, 15 s at 55°C and 30 s at 72°C. A final elongation step of 5 minutes at 72 °C was applied. PCR fragments were purified using QIAquick PCR Purification kit (Qiagen) and sequenced in an ABI 3130xl Genetic Analyzer instrument (Thermo Fisher Scientific) using the BigDye v3.1Terminator Cycle Sequencing Kits. Electropherograms underwent examination using the Phred algorithm within a tool hosted on the webpage http://www.biomol.unb.br/phph/. Sequences were compared against the NCBI nucleotide database utilizing the Blastn algorithm (25).

### Genome sequencing, assembly and annotation

After confirming the isolates as *Sarocladium*, genome sequencing was performed using one microgram of DNA per isolate for paired-end sequencing. DNA was fragmented via sonication, followed by size selection at 500 base pairs. Library preparation followed the Kapa Biosystems kit protocol (Kapa Biosystems, Woburn, MA), incorporating an 8-base pair index for multiplexing. Library quality was assessed using quantitative PCR (qPCR) on a 7900HT system (Life Technologies Corporation, Carlsbad, CA) with a Kapa library quantification kit (Kapa, Woburn, MA). Sequencing was carried out in an Illumina MiSeq instrument under high-output mode, generating 2 × 300 base pair paired-end reads. Demultiplexing was performed using in-house tools.

We assembled and annotated the two resulting genomes, *S. spinificis* CA16 and CA18. We used the AAFTF/0.4.1 pipeline (26) for assembly, and the fungal genome annotation pipeline, funannotate v1.8 (Palmer and Stajich 2020), to annotate them. We also identified and masked repetitive DNA elements using TANTAN (27). To do gene prediction, we used the funannotate “predict” command, incorporating evidence-based, and *ab initio* (structure-based) prediction pathways. To train the *ab initio* predictors Augustus (28), GlimmerHMM (29), and SNAP (30), we retrieved conserved gene models from the sordariomycetes_odb10 BUSCO database (31). Additionally, we also used GeneMark-ES was with the option tailored for fungal genomes to further enhance ORF prediction and improve the accuracy of gene model annotation (32). We inferred a weighted consensus gene structure via EVidenceModeler (33), with a weight of 2 assigned to Augustus HiQ models and a weight of 1 for others. We also predicted tRNAs using tRNAscan-SE (34). We filtered out genes with a length less than 50 amino acids and those identified as transposable elements for all downstream analyses.

We investigated gene family expansions by comparing the six available *Sarocladium* genomes against multiple functional annotation databases. Protein function was assigned using eggNOG 5.0 (35), Pfam v36.0 (36), and InterProScan v96.0 (37). Additionally, specific functional categories, including carbohydrate-degrading enzymes (CAZy/dbCAN v11.0 (38)), proteases (MEROPS v12.0 (39)), and secreted proteins (40), were analyzed to gain further insights into metabolic capabilities and potential pathogenicity factors. We also did Gene Ontology (GO) analyses with WHAT TOOL (41). Finally, we predicted biosynthetic gene clusters (BGCs) were using fungiSMASH, a fungal-genome-specific version of antiSMASH v6.0 (42), in the two *Sarocladium* genomes.

### Phylogenetic trees

Next, we explored the phylogenetic relationships of the two sequenced isolates with other *Sarocladium* isolates. We used two approaches, Multi-Locus Sequencing Typing (MLST) and phylogenomic analysis. We described the protocols to generate input data, and run the phylogenetic analyses as follows.

*<u>i)</u>* <u>MLST-based tree</u>: Most fungal data available for phylogenetics exists in the form of sequencing of a few loci. In particular, the loci LSU, ITS, TEF, RPB2 and ACT are often used in fungal phylogenetics (4, 43, 44). We retrieved these five genetic markers from the CA16 and CA18 genomes using Blastn analysis and aligned to other 54 related taxa (Supplementary Table 1) using the MAFFT online server (45). It is worth noting that some isolates of *S. spinificis* were previously classified as distinct species before phylogenetic analyses clarified their taxonomic placement (4). Manual curation was performed and phylogenetically-informative sites from each genetic maker were collected using the ClipKIT v1.3.0 function smart-gap (46). Phylogenetic trees were inferred using the maximum likelihood (ML) method implemented in IQ-TREE2 (47). The nucleotide substitution model was automatically selected using the *-m TEST* function, which determines the best-fitting model based on the Akaike Information Criterion (AICX - (48). To assess branch support, we performed ultrafast bootstrap approximation (UFBoot - (49)) and approximate likelihood ratio tests (aLRT - (50)). The resulting tree was visualized using FigTree software v 1.4 (http://tree.bio.ed.ac.uk/software/figtree/). We used *Parasarocladium wereldwijsianum* strain NL19095011 as an outgroup to root the tree.
*<u>ii)</u>* <u>Phylogenomic tree</u>: To generate phylogenetic trees, we used the PHYling pipeline (https://github.com/stajichlab/PHYling), which extracts phylogenetically conserved markers and constructs species trees from annotated genomes. This approach was complemented by a genome-wide phylogenetic analysis with a smaller taxonomic representation, providing an additional perspective on evolutionary relationships among the studied taxa. We used the set of Benchmarking Universal Single-Copy Orthologs (BUSCO) listed in the fungi_odb10 database (31). We retrieve the protein models in five previously published *Sarocladium*: *S. implicatum* TR *-* GCA_021176775.1, *S. kiliense* ZJ-1 - GCA_030734395.1, *S. strictum* F4-1 - GCA_030435815.1*, S. oryzae* JCM 12450 - GCA_001972265.1 and *S. brachiariae* HND5- GCA_008271525.1. We used the hmmsearch tool from HMMER v3.3.2 (51) to retrieve BUSCO orthologous genomic loci. We generated protein alignments using the hmmbuild function of HMMER v3.3.2 (51) and the ClipKIT v 1.3.0 tool (46), using the smartgap function, was utilized to eliminate spurious positions. *Tolypocladium ophioglossoides* strain CBS100239 was used as outgroup (52). We applied genome-wide phylogenetic analysis using the genealogical concordance approach on IQ-TREE 2 software (47). We used the -m TEST function to infer the best nucleotide and protein alignment models. To asses branch support, we calculated 1,000 ultrafast bootstraps (49), the approximate likelihood-ratio test (aLRT) (50), and approximate Bayes test (abayes) for each node (53). We visualized the resulting phylogenetic trees were visualized using the FigTree v1.4.4 software (https://tree.bio.ed.ac.uk/software/figtree/).

### Comparative genomics

We used the genome data of CA16 and CA18, and the previously sequenced *Sarocladium* genomes to identify gene family expansions. We next run the funannotate v1.8 compare function in the six *Sarocladium* species, to look for genomic trends of the annotated protein categories (see Genome sequencing, assembly and annotation section). Heat maps were produced from those uneven counts of protein families or classes with standard deviation greater than 1.0. We visualized shared and unique orthologous clusters in *Sarocladium* using Orthoveen3 (54).

## RESULTS

### Clinical cases

We identified and recovered two *Sarocladium* sp. isolates, CA16 and CA18, at the Fungus Testing Laboratory at the University of Texas Health San Antonio, TX. The strain CA16 was obtained from a patient presenting with a persistent ulcerative lesion on the lower lip, characterized by chronic inflammation. The lesion had an indurated border and displayed progressive erosion, raising suspicion of a fungal etiology. Similarly, isolate CA18 was recovered from a chronic, non-healing wound on the left lower leg of a different patient, also from Madera, California. The wound initially developed following minor trauma but failed to heal despite standard wound care and antibiotic therapy. Biopsy and mycological analysis led to the identification of a filamentous fungi as the etiological agent. Coccidioidomycosis was ruled out as qPCR-based assays using a *Coccidioides*-specific test yielded negative results.

### Morphological and molecular characterization of CA16 and CA18 strains

Both CA16 and CA18 strains grew at 24°C on PDA, MEA and OA agar showing similar micromorphology with little phenotypic variability (Figure 1A-C, Figure S1). On both temperatures the strains grew as filamentous forms. Colonies circular, regular, flat, whitish or creamy buff, sometimes radially or slightly sulcate. In all media, yellow to orange pigment was produced. However, at 37 °C colonies produced no pigmentation and grew slower (Figure 2). The macrocolonies of CA16 (Figures 2A-B) and CA18 (Figures 2C-D) grown on 2X-GYE media at 24°C for 15 days measured 44 mm and 36 mm, respectively. Colonies were irregular and wrinkled, with a pale orange center, white margins, and an orange reverse, while both strains produced a soluble yellow pigment diffused in the agar (Figure 1A-C, Figure S1). At 37°C, CA16 (Figures 2E-F) and CA18 (Figures 2G-H) reached 42 mm and 24 mm after 15 days of growth respectively, forming irregular, cottony white colonies with a pale to yellowish reverse (Figure S1). Microscopically, hyphae septate, branching, smooth and hyaline, some arranged in rope-like appearance; conidiophores phialidic, undifferentiated; phialides erect, mostly solitary, rarely branching, cylindrical, with inconspicuous collerates, producing conidia in slimy-head shaped; conidia are cylindrical or ellipsoidal measuring from 2µm to 7µm (Figure 3). Both isolates showed moderate resistance to antifungals. Table 1 shows the minimum inhibitory concentrations (MICs) for the two S. *spinificis* isolates.

**Figure 1.**
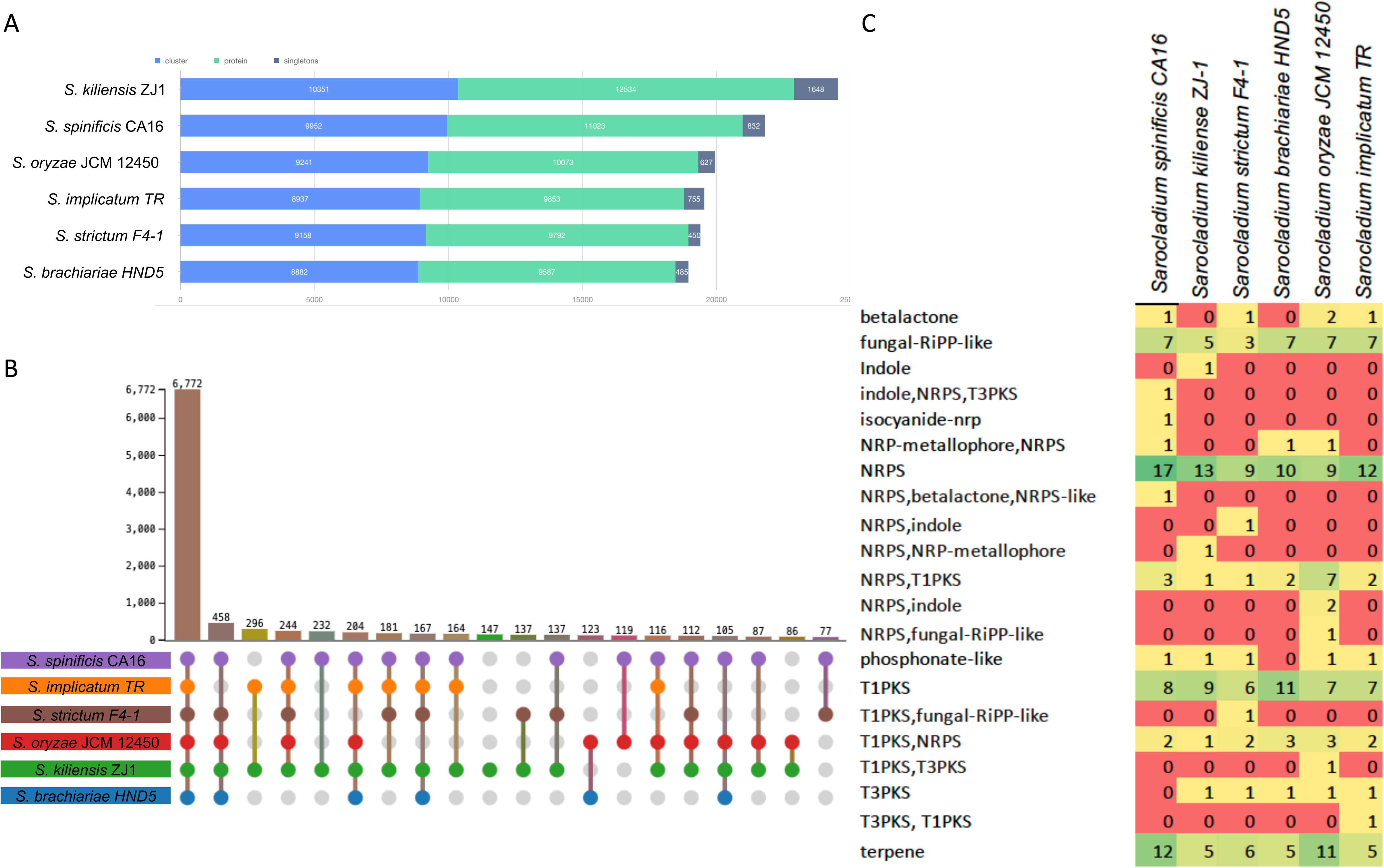
Growth and micromorphology of *Sarocladium spinificis* strains CA16 and CA18. Both strains exhibited growth at 24°C on PDA (A), MEA (B), and OTA (C) agar, displaying similar micromorphology with minor variations in colony morphology. Microscopically, hyphae are septate, branching, smooth, and hyaline, with some forming a rope-like appearance. Conidiophores are phialidic and undifferentiated, with erect, mostly solitary phialides that are rarely branched, cylindrical, and possess inconspicuous collarettes. Conidia are produced in slimy-head formations, appearing cylindrical to ellipsoidal and measuring 2–7 µm in length (D-H)

**Figure 2.**
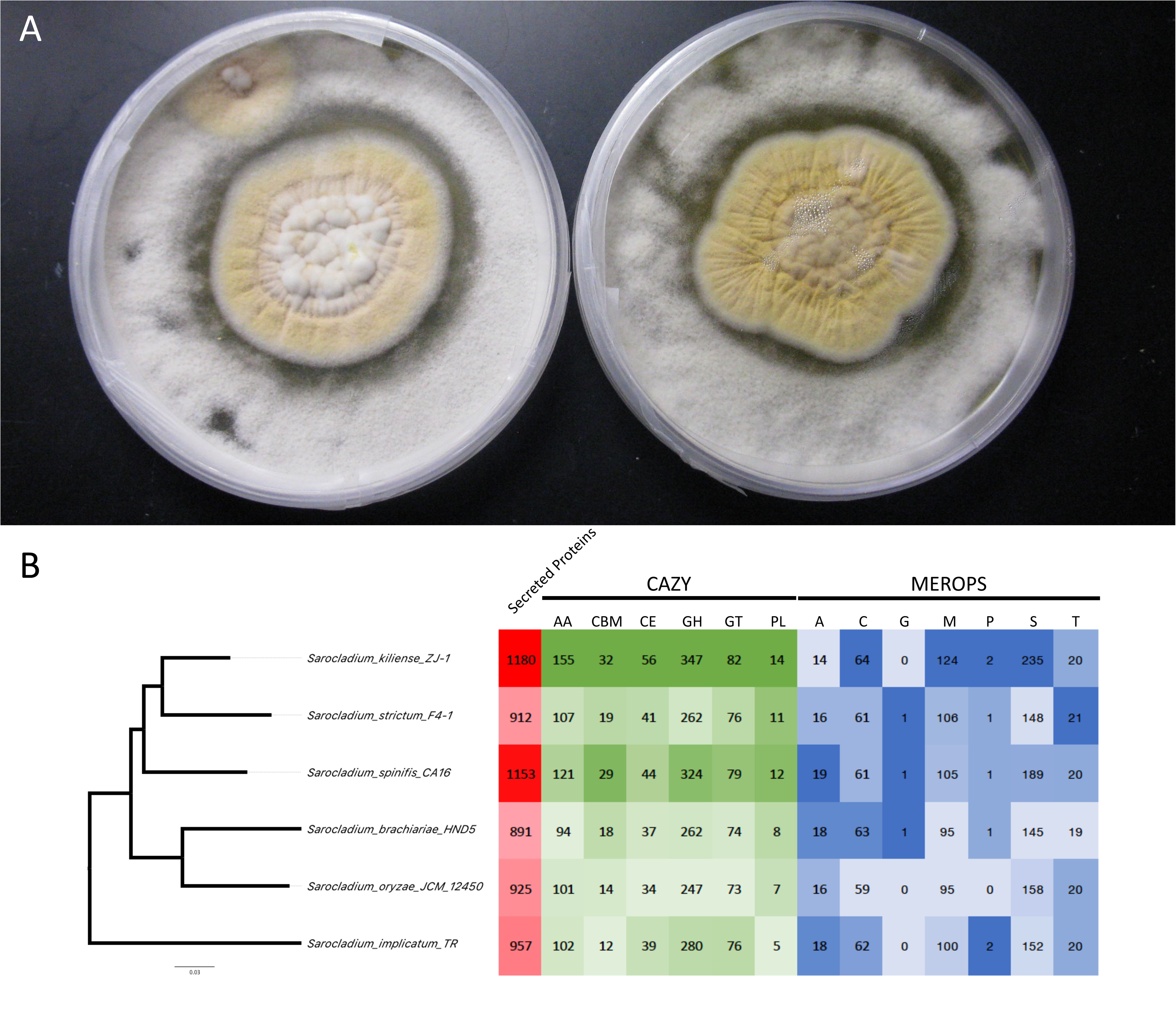
Growth characteristics of *Sarocladium spinificis* strains CA16 and CA18 at 24°C and 37°C. (A–B) Macrocolonies of CA16 and (C–D) CA18 grown on 2X-GYE agar at 24°C for 15 days, measuring 44 mm and 36 mm, respectively. Colonies were irregular and wrinkled, with a pale orange center, white margins, and an orange reverse, while both strains produced a soluble yellow pigment diffused in the agar. (E–F) CA16 and (G–H) CA18 grown at 37°C for 15 days, reaching 42 mm and 24 mm, respectively. Colonies at this temperature were irregular, cottony white, with a pale to yellowish reverse, and showed reduced growth with no pigmentation.

**Figure 3.**
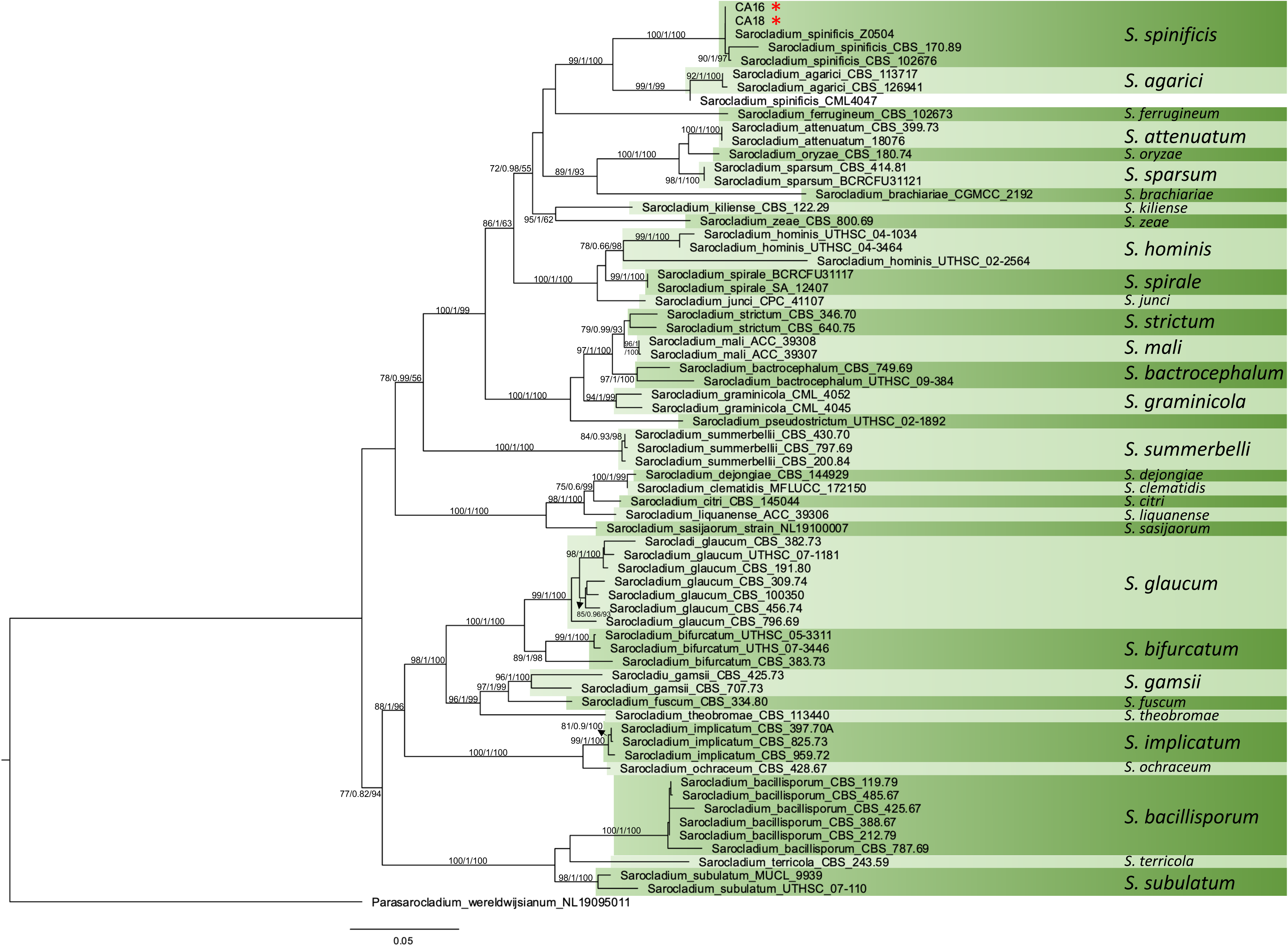
MLST-based phylogenetic tree of *Sarocladium spinificis*. Maximum likelihood (ML) phylogenetic tree inferred from concatenated sequences of five loci (LSU, ITS, TEF, RPB2, and ACT) retrieved from *S. spinificis* strains CA16 and CA18, along with 54 related taxa (Supplementary Table 1). Phylogenetic analyses were performed using IQ-TREE2, with model selection determined by the Akaike Information Criterion (AIC). Branch support was assessed using ultrafast bootstrap approximation (UFBoot) and approximate likelihood ratio tests (aLRT). The tree was visualized in FigTree v1.4, with *Parasarocladium wereldwijsianum* strain NL19095011 used as the outgroup for rooting. Branch lengths in the ML tree are proportional to the number of nucleotide substitutions per site, reflecting genetic divergence between taxa.

**Table 1.** Minimum inhibitory concentrations (MICs) of four antifungal agents for *Sarocladium spinificis* isolates CA16 and CA18. All measurements are in μg/ml.

|  | CA16 | CA18 |
| --- | --- | --- |
| Amphotericin B | 0.125 | 0.06 |
| Fluconazole | 16 | 16 |
| Itraconazole | 0.5 | 1.0 |
| Voriconazole | 0.25 | 0.06 |

Next, we amplified and sequenced the universal fungi barcode rDNA locus ITS for both CA16 (MF784843.1) and CA18 (MF784844.1) strains. Blastn analyses reveled that those two isolated were 99.82% identical to *S. spinificis* strain Y. H. Yeh I0317 (MK336486.1), isolated from a Beach Morning Glory (*Ipomoea pes-caprae*) plant in Taiwan. The maximum likelihood MLST-based phylogenetic tree indicated that both CA16 and CA18 clustered with three isolates of *S. spinificis*. The branch leading to these five isolates had high levels of support (bootstrap and aLRT; Figure 3). Two *S. spinificis* strains, CBS 170.89 and CBS 102676, form a discrete dyad, but we have no power to resolve the relationships between the other three strains of the group (CA16, CA18, and Z0504). This clade is sister to a clade formed by with *Sarocladium agarici* and a sixth isolate of *S. spinificis* (CML4047). The sister relationship *spinificis* and agarici is concordant with previous results (4), but the position of CML4047 latter isolate is puzzling.

We also used a phylogenomic approach based on 758 protein markers to infer the phylogenetic relationships between *Sarocladium* lineages. This analysis included fewer isolates but had more phylogenetic power than out MLST analysis. *S. spinificis* CA16 shares a common ancestor with *Sarocladium kiliense* ZJ-1 and *Sarocladium strictum* F4-1 (Figure 4A). This is in disagreement with the former MLST analysis due discrepancy in terms of number of taxa analyzed (Figure 3). Unfortunately, there were only six *Sarocladium* species represented by whole genomes at the time of the analysis, however both MLST and phylogenomic analyses are complimentary.

**Figure 4.**
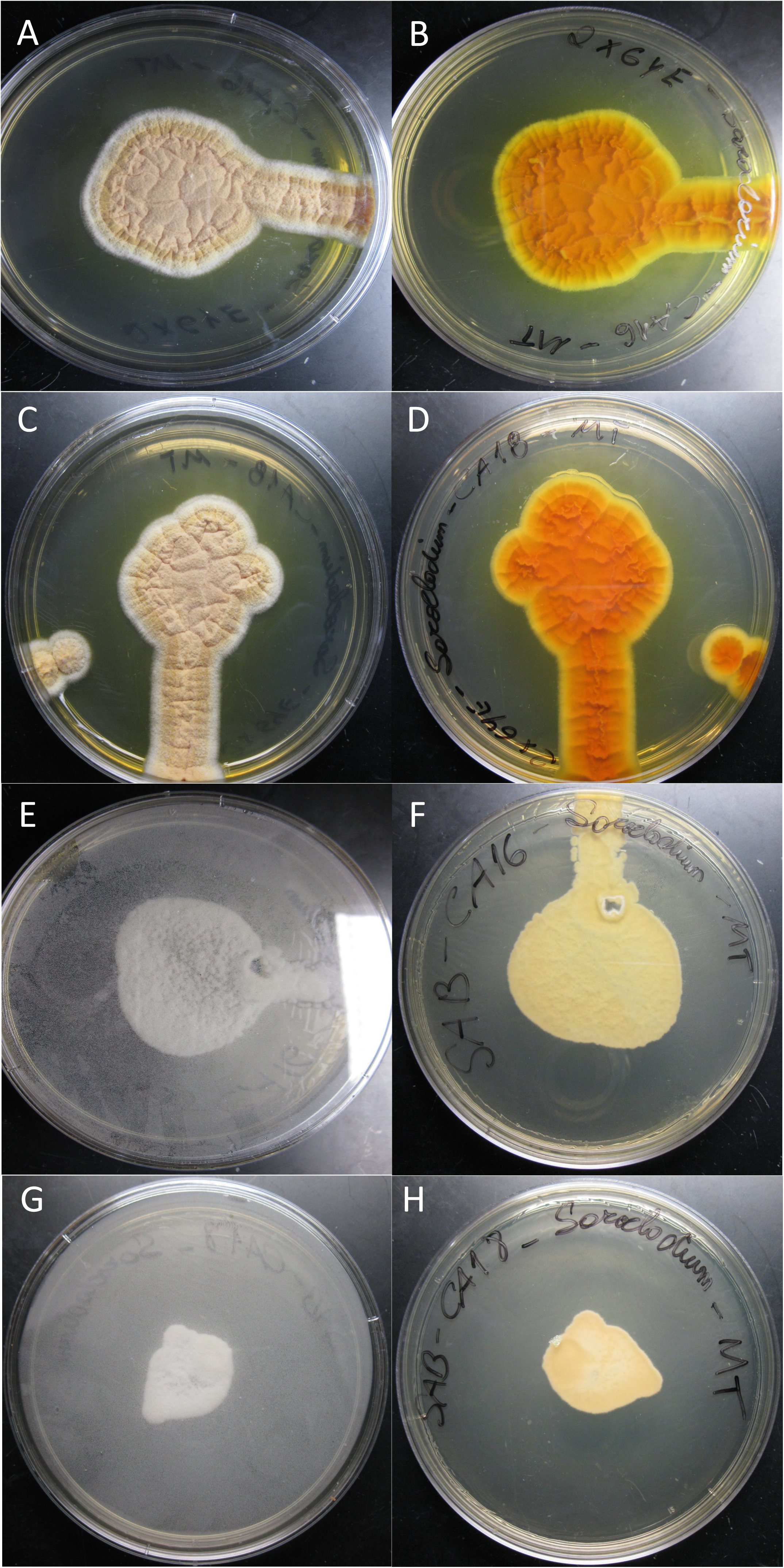
Antagonistic interactions and phylogenomic relationships of *Sarocladium spinificis*. (A) After 10 days of growth, a distinct halo of inhibition was observed between the margin of *S. spinificis* CA16 or CA18 and *Coccidioides posadasii* NR-166, suggesting that *S. spinificis* is capable of inhibiting or restricting the growth of *C. posadasii*. (B) Phylogenomic relationships among *Sarocladium* species were inferred using 758 conserved protein markers extracted via the PHYling pipeline. This approach, which provides greater phylogenetic resolution than MLST analysis, confirmed that *S. spinificis* CA16 shares a common ancestor with *S. kiliense* ZJ-1 and *S. strictum* F4-1. Heat maps represent functional annotations for CAZy/dbCAN v11.0 (carbohydrate-active enzymes), MEROPS v12.0 (proteases), and secreted proteins, highlighting key metabolic differences among *Sarocladium* species.

### Sarocladium spinificis CA16 inhibits the grow of Coccidioides posadasii

Both patients that were infected by the strains CA16 and CA18 were initially diagnosed with coccidioidomycosis. ACCUPROBE testing confirmed *Coccidioides* sp. in the cultures but morphological and qPCR analyzes ruled out this possibility. Nevertheless, *S. spinificis* can also grow at 37°C, show macromorphological characteristics that may resemble, but do not entirely match, those of *Coccidioides*, which can cause diagnostic confusion. These factors make it challenging to determine whether it was a co-infection, a misdiagnosed case of coccidioidomycosis, or if *S. spinificis* is merely a thermotolerant contaminant. To understand how those microorganisms interact, we co-cultivated both fungi in the same petri dish. After 10 days of growth, a distinct halo of inhibition was observed between the margin of *S. spinificis* CA16 or CA18 and *C. posadasii* NR-166, suggesting that *S. spinificis* is capable of inhibiting, or at least restricting, the growth of *C. posadasii* (Figure 4A).

### Whole genome analysis and annotation

The genomes of *S. spinificis* CA16 or CA18 were assembled into 414 and 368 contigs and yielded 33.61Mb and 33.73 Mb respectively (Table 2). Desite their fragmentation, the genomes are largely completed according to BUSCO analysis (CA 16 – 98% and CA18 – 98.2% - Table 2). The content of repetitive DNA in the *S. spinificis* genome was lower (CA16 – 2.34% and CA18 – 2.47%) than that of other *Sarocladium* genomes (Table 2), such as *S. kiliense* (7.04%) and *S. strictum* (8.49%).

**Table 2.**
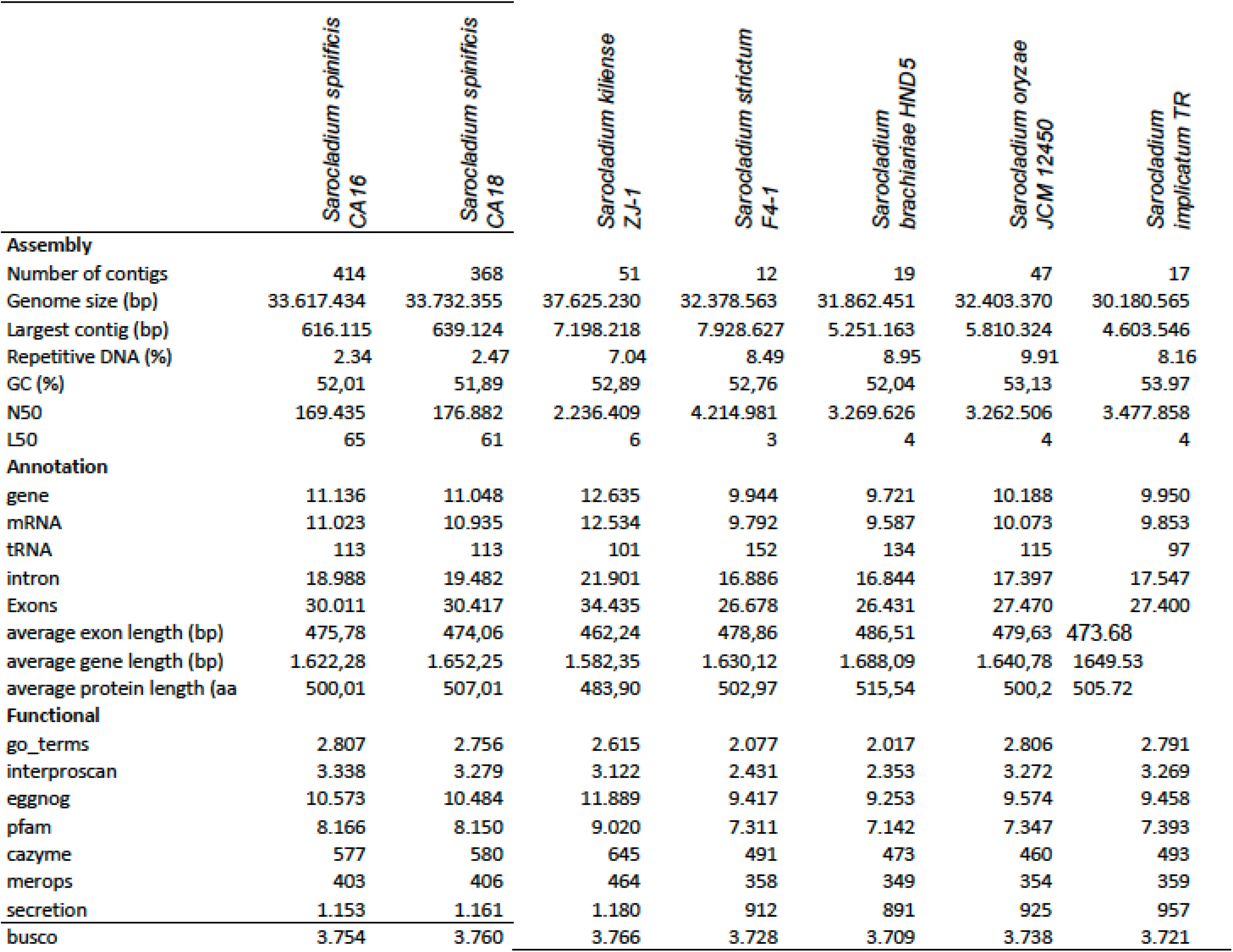
Summary of whole-genome assemblies and annotation of the Sarocladium species

In terms of number of protein-coding genes, *S. kiliense* harbors 12,635 while *S. spinificis* CA16 or CA18 strains has 11,023 and 10,935 respectively while the other *Sarocladium* species has a lower number of predicted genes (Table 2). On a similar vein, *S. kiliense* has 10,351 orthologous clusters and 1,648 singletons, *S. spinificis* has 9,952 orthologous clusters and 832 singletons, and the remaining *Sarocladium* species have fewer orthologous clusters and singletons (Figure 5A).

**Figure 5.**
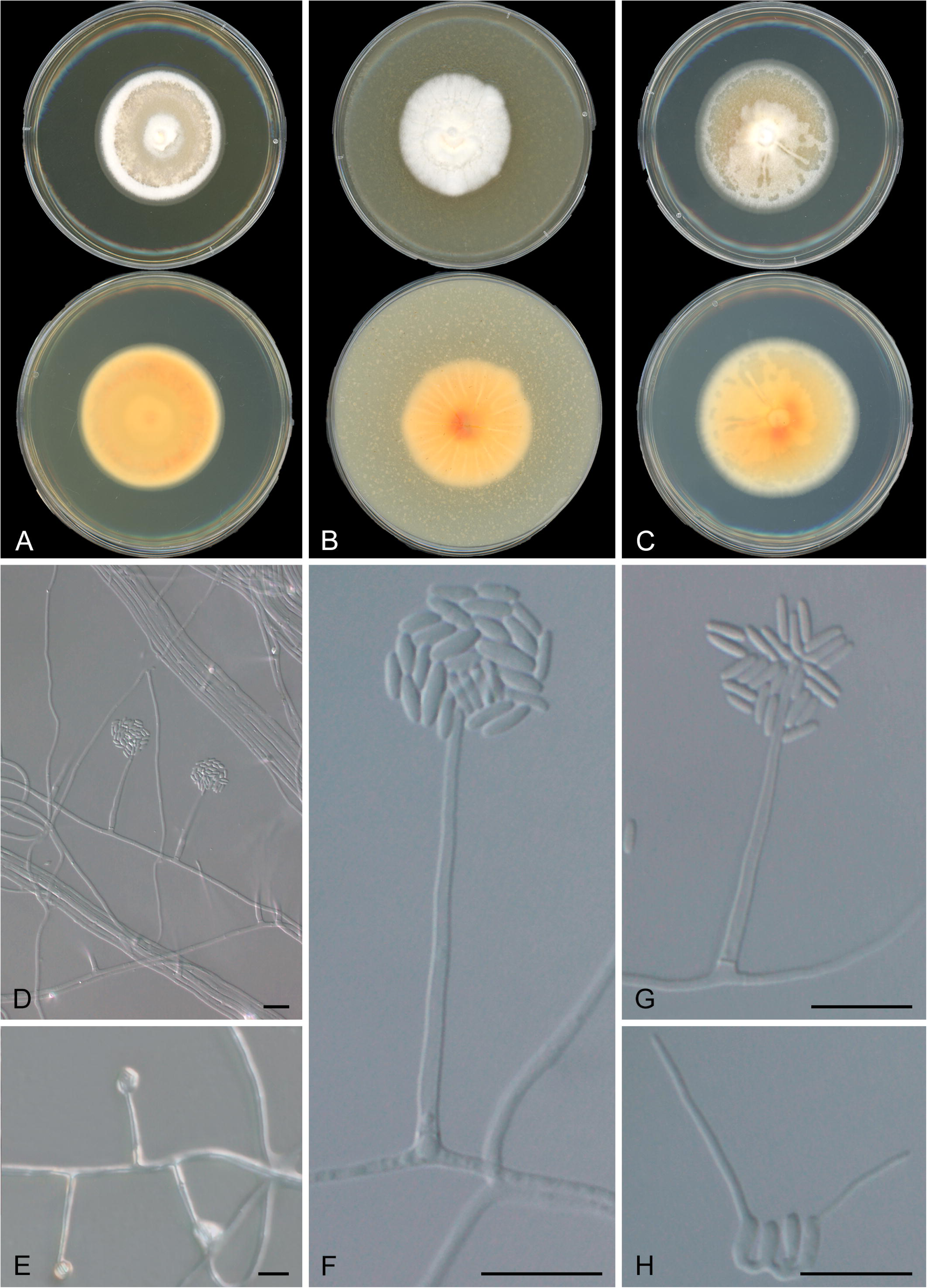
Orthologous cluster analysis was performed using OrthoVenn3, visualizing shared and unique protein clusters among *Sarocladium* genomes. Venn diagrams depict the distribution of conserved and species-specific clusters, shedding light on genomic diversity and evolutionary relationships (A-B). Secondary metabolite biosynthetic potential was assessed using antiSMASH, revealing key differences in the presence and composition of biosynthetic gene clusters involved in metabolite production across *Sarocladium* species (C).

In total, we found 6,772 shared orthologous clusters across the six *Sarocladium* species (Figure 5A). We also found 29 unique orthologous clusters for *S. spinificis* and according to GO analysis those are related to lipid and carbohydrate metabolism, cellular aromatic compound metabolic process (I.E. terpene) and transport (Biological Process) (Table S1). The only molecular Function categories that were overrepresented were, monooxygenase and oxidoreductase activities. Genes involved response to acidic pH (biological process GO:0010447) was significant enriched in S. *spinificis* compared to the other species (p-value = 0.0014).

We found similarities in protein family evolution between *S. kiliense* and *S. spinificis* which differentiated them from other *Saracocladium* genomes. We have found an increase in eggNOG terms, Pfam, CAZymes, MEROPS (peptidases), and secreted proteins in *S. kiliense* and *S. spinificis* compared to *S. strictum*, *S. brachiariae* and *S. oryzae* which might reflect in their different ecological roles and physiology (Figure 4A, Table 2). We found 232 orthologous clusters shared that have undergone gene expansion in both *S. kiliense* and *S. spinificis*, but not in other species. The genomic similarities between these two species are noteworthy because S. *kiliense* and and *S. spinificis* have been implicated in opportunistic infections in humans and both can grow at 37°C, a characteristic common to most fungal human pathogens.

Our analysis identified expanded CAZy gene families in *Sarocladium kiliense* and *Sarocladium spinificis*. These included eight glycoside hydrolase (GH) families (GH18, GH43, GH16, GH5, GH78, GH35, GH109, CBM91), a monooxygenase family (AA19), a glucose-methanol-choline (GMC) oxidoreductase family (AA3), and four peptidase families (SX09, S10, S33, M14B), all of which were enriched in their genomes (Figure 4B). MEROPS M38 protein family, a peptidase family of isoaspartyl dipeptidases is highly expanded in *S. kiliense* (Figure 4B) and members of this family are known to play a role in virulence and pathogenesis in different fungal pathogens (55, 56).

We found a similar trend in transcription factors, as we found gene family expansions of genes with TF domains in *S. kiliense* and *S. spinificis*. IPR007219, a fungal-specific TF domain was enriched in the *S. kiliense* and *S. spinificis* genomes compared to the other *Sarocladium* species. Two TF domains showed expansions in plant pathogens. IPR000679, a GATA zinc finger domain, was enriched in *S. implicatum*; and IPR004827, a basic region leucine zipper 2 domain and IPR004827, bZIP TF 1 are enriched in *S. oryzae* (Figure S2).

We annotated 21 secondary metabolite gene groups in the six *Sarocladium* genomes (Figure 5C), and found notable differences. The indole-NRPS-T3PKS, NRPS-betalactone- NRPS-like, and isocyanide-nrp clusters are uniquely present in *S. spinificis* CA16 genomes, while the T3PKS cluster is absent compared to other *Sarocladium* species. We also observed an overrepresentation of NPRS and terpene clusters in the *S. spinificis* genome in agreement with GO enrichment analysis (Figure 5C).

## DISCUSSION

In this study, we report that *Sarocladium* spinificis can be associated with human infections concomitantly with *Coccidioides*. We characterized two *Sarocladium* strains, CA16 and CA18, both recovered from patients suspected of having coccidioidomycosis (Figure 1, Figure S1), using morphological and molecular approaches. Molecular assays ruled out *Coccidioides* species and confirmed that the strains belonged to the genus *Sarocladium*. Identification was further refined through rDNA ITS locus amplification and sequencing, supported by phylogenetic analysis of multiple genetic loci, including whole genome analyses. These analyses revealed a close relationship between CA16, CA18, and isolates of *Sarocladium spinificis* derived from plant-derived sources. Previous reports have also identified *Parengyodontium* species in cases suspected of coccidioidomycosis, expanding the range of fungi involved in such clinical presentations (57). The classification of these fungi as contaminants, opportunistic pathogens, or primary fungal pathogens remains an open question for further investigation. We also report that the two *S. spinificis* are able to outcompete or inhibit the growth of *Coccidioides*. Our findings contribute to the understanding of *S. spinificis* in two key aspects: as a human commensal capable with the potential of becoming an opportunistic pathogen, and as a fungus that can suppress the growth of a major human pathogen. We discuss these two facets in the following sections.

*Sarocladium* species are a recently recognized human pathogen and little to nothing is known regarding their prevalence, or the molecular underpinnings behind virulence in mammals. *Sarocladium kiliense* caused an outbreak of bloodstream infections in Chile and Colombia associated a batch of contaminated antinausea medication in immunosuppressed (17). All isolates were closely related which confirm a single source (17). It is worth noting that species of *Acremonium*, the former genus of *S. kiliense*, have also been implicated in human disease (reviewed in (8)). Although our *S. spinificis* isolates were recovered from human patients, we cannot conclusively determine whether this species causes disease in humans or other mammals. Nonetheless, S. *spinificis*’ growth at 37°C is significant, as it matches human body temperature, suggesting that these fungi can survive and potentially thrive in a human host. The ability of these fungi to grow at this temperature coupled with effects of climate change indicates their potential to emerge as pathogens causing subcutaneous mycotic infections (58). We also report antifungal resistance in S. *spinificis*, which follows observations in *S. kiliense* (59). Experimental infections in mice or human cell lines are necessary to establish whether *S. spinificis*, like *S. kiliense*, has the potential to become a human health threat.

Our findings also provide new insights into the evolution of *Sarocladium*. Phylogenetic analysis of multiple loci confirmed that CA16 and CA18 are closely related to *S. spinificis*, but not to *S. kiliense* (Figure 3-4). This raises the question of whether other *Sarocladium* species can also cause disease—or at least persist as human commensals—or if virulence mechanisms evolved independently in these species. Comparative genomics revealed that *S. spinificis* and *S. kiliense* have a higher number of secreted proteins, CAZymes, and peptidases, which may contribute to their ability to adapt to diverse ecological niches and their potential pathogenicity (Figure 4). Additionally, the presence and diversity of monooxygenases and oxidoreductases suggest metabolic versatility. These traits appear to have evolved in parallel, indicating similar selective pressures (Table S1). These findings have broader implications. *Sarocladium* was previously classified within the genus *Acremonium* (1, 8). a genus that harbors other human pathogens (60). Determining what are the evolutionary mechanisms that have driven the evolution of virulence in the *Acrenomiun/Sarocladium* clade will require the resolution of the phylogenetic relationships between the species in the group using genome wide markers.

Our co-cultivation experiment with *S. spinificis* and *Coccidioides posadasii* was particularly striking, as CA16 and CA18 originated from suspected coccidioidomycosis cases. Both strains not only inhibited but also significantly outcompeted *C. posadasii,* suggesting potential competitive interactions or the production of antifungal compounds. Similar antagnositic interactions have been observed in other members of the order Hypocreales, such as *Trichoderma* (61, 62), *Acremonium* (63, 64), and *Beauveria* (65, 66), which produce antifungal metabolites and enzymes. A different species of *Sarocladium*, *S. brachiariae* produces volatile compounds able to inhibit the broth of *Fusarium oxysporum* f. sp. *cubense* (67). These findings highlight the ecological and clinical significance of *S. spinificis* and its potential role in shaping microbial communities and disease dynamics. Our study provides a comprehensive characterization of CA16 and CA18, offering new insights into their genome, ecological interactions, and phylogenetic relationships. By expanding our understanding of *Sarocladium* diversity and pathogenic potential, this work lays the foundation for future research on its role in both environmental and clinical contexts, as well as its potential applications in biotechnology and drug discovery.

---

**Figure S1.** Filamentous growth of *Sarocladium spinificis* CA16 and CA18 strains. Fungal cultures were grown at 24°C on Potato Dextrose Agar (CA16, A-B and CA18, C-D), Oatmeal Agar (CA16, E-F and CA18, G-H) and Malt Extract Agar (CA16, I-J and CA18, L-M showing similar macromorphology with little phenotypic variability. Colonies circular, regular, flat, whitish or creamy buff, sometimes radially or slightly sulcate. In all media, yellow to orange pigment was produced.

**Figure S2.** Expansion of Transcription Factor Gene Families in *Sarocladium* Species. We identified transcription factor (TF) gene family expansions in *Sarocladium kiliense* and *Sarocladium spinificis* using genome data from CA16, CA18, and previously sequenced *Sarocladium* genomes. To analyze genomic trends in annotated TF categories, we employed the Funannotate v1.8 compare function across six *Sarocladium* species. Our analysis revealed that IPR007219, a fungal-specific TF domain, was significantly enriched in *S. kiliense* and *S. spinificis* compared to other *Sarocladium* species. Additionally, we observed expansions of transcription factors associated with plant pathogens. Specifically, IPR000679 (GATA zinc finger domain) was enriched in *S. implicatum*, while IPR004827 (basic region leucine zipper 2 domain, bZIP TF 1) was significantly enriched in *S. oryzae*

