## Supplementary figures and images for "A Novel *Sarocladium spinificis* Strain Suppresses *Coccidioides posadasii* Growth: Morphological and Genetic Perspectives"

### Figure S1

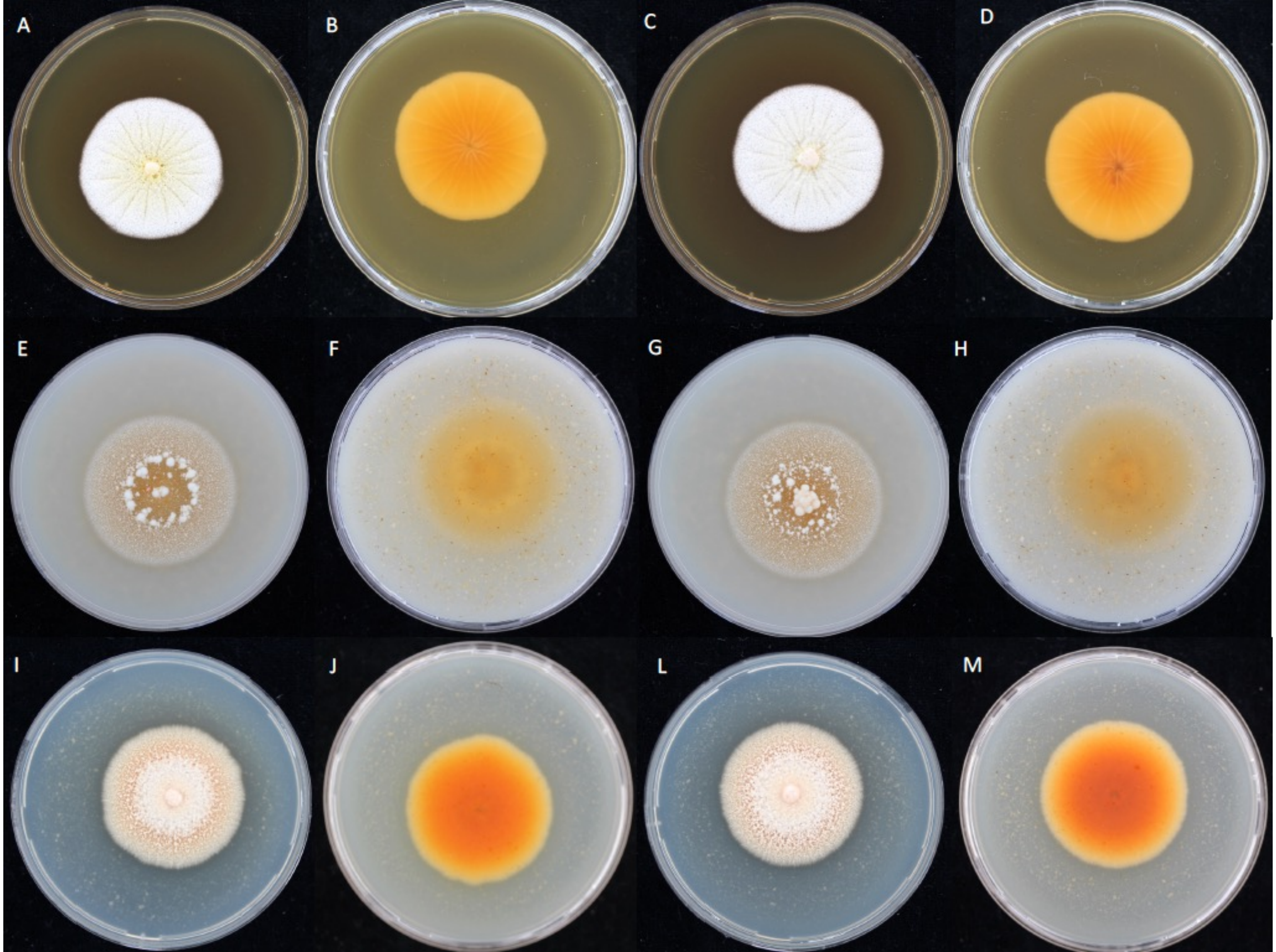

### Figure S2

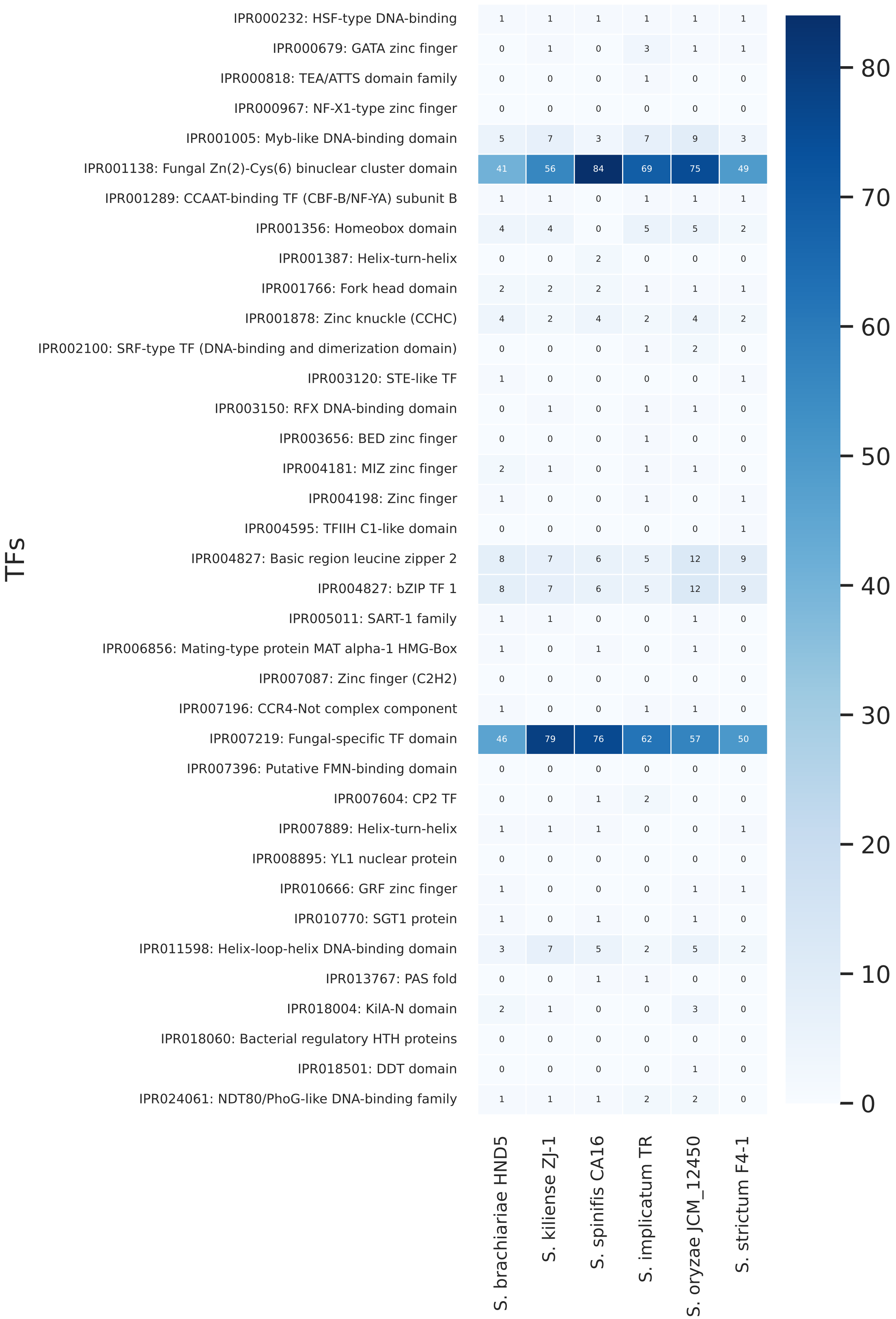
